# Survival of the salient: Emotion rescues otherwise forgettable memories via neural reactivation and post-encoding hippocampal connectivity

**DOI:** 10.1101/2020.07.07.192252

**Authors:** David Clewett, Joseph Dunsmoor, Shelby Bachman, Elizabeth Phelps, Lila Davachi

**Author notes:** To whom correspondence should be addressed: Department of Psychology, Columbia University, Schermerhorn Hall #406, 1190 Amsterdam Ave, New York, NY 10027.

## Abstract

Emotion’s selective effects on memory go beyond the simple enhancement of threatening or rewarding stimuli. They can also rescue otherwise forgettable memories that share overlapping features. Here, we use functional magnetic resonance imaging (fMRI) to examine the brain mechanisms that support this retrograde memory enhancement. In a two-phase incidental encoding paradigm, participants first view images of neutral tools and animals. During Phase 1, these images are intermixed with neutral scenes, which provides a unique ‘context tag’ for this specific phase of encoding. A few minutes later, during Phase 2, new pictures from one category are paired with a mild shock (fear-conditioned stimulus; CS+), while pictures from the other category are not shocked. fMRI analyses reveal that, across participants, retroactive memory benefits for Phase 1 CS+ items are associated with greater phasic reinstatement of the prior mental context during Phase 2 CS+ items. We also see that greater VTA/SN activation during Phase 2 CS+ items relates to this retroactive memory enhancement, suggesting that emotion promotes both the encoding and ongoing consolidation of overlapping representations. Additionally, we find that emotional experience-dependent changes in post-encoding hippocampal functional coupling with CS+ category-selective cortex relate to the magnitude of the retroactive memory effect. These hippocampal connectivity patterns also mediate the relationship between dopaminergic emotional encoding effects and across-participant variability in the retroactive memory benefit. Collectively, our findings suggest that an interplay between online and offline brain mechanisms may enable emotion to preserve seemingly mundane memories that become significant in the future.

## Introduction

Years of research show that emotionally arousing stimuli are preferentially processed and remembered, often at the expense of processing temporally and/or spatially adjacent neutral information (Kensinger et al., 2007; LaBar & Cabeza, 2006; Mather & Sutherland, 2011; Sakaki et al., 2014). In the real world, however, we don’t always know that something is or, indeed, *was* relevant until later on. Thus, while some occurrences might seem inconsequential in the moment, an adaptive memory system should retain some seemingly mundane information, at least temporarily, in case this information gains significance in the future. Eating a bland sandwich at a new restaurant, for example, will likely form a weak memory trace that fades quickly over time. But if you experienced food poisoning soon after, it would be important to preserve that recent memory so that you can remember to avoid that restaurant in the future. The goal of the present study was to acquire a deeper understanding of how emotionally-significant events prioritize memory for only certain aspects of a recent, seemingly mundane experience.

Experiencing something arousing after an event can strongly influence memory consolidation for recently studied words or images. This effect has been demonstrated robustly with stressors (Andreano & Cahill, 2006; McCullough & Yonelinas, 2013; Sazma, McCullough, et al., 2019), norepinephrine (Southwick et al., 2002), acute exercise (Nielson et al., 1996), reward (Braun et al., 2018; Murayama & Kitagami, 2014; Patil et al., 2017), novelty (Ballarini et al., 2009), and emotional videos (Nielson & Powless, 2007; Nielson et al., 2005). Similar general memory enhancements have been reported in rodents when weak learning is followed by stimulating exploration of a novel open field (Moncada et al., 2011) or epinephrine injections (McGaugh & Roozendaal, 2002). Across many emotion-cognition studies, post-encoding arousal can also selectively enhance memory for emotional over neutral information (Cahill & Alkire, 2003; Goldfarb et al., 2019; Sazma, Shields, et al., 2019; Smeets et al., 2007), suggesting that arousal may privilege the consolidation of motivationally-relevant events even after those experiences have transpired.

But this body of work does not tell the whole story of how emotional events impact memory consolidation. While less common, some studies show that post-encoding arousal can enhance memories for a subset of preceding neutral stimuli (Liu et al., 2008; Preuss & Wolf, 2009; Sazma, Shields, et al., 2019; Smeets et al., 2007). How, then, do we predict which neutral memories will be enhanced by post-encoding emotional arousal? Recent findings in humans demonstrate that emotional learning selectively and retroactively enhances memory for conceptually-related neutral information (Dunsmoor et al., 2015; Patil et al., 2017). Conceptual overlap between an arousal source and the to-be-learned information thereby appears to be a key factor in determining which neutral representations are preserved by a subsequent emotionally arousing context. While intriguing, the neural processes that support this selective retroactive memory benefit are unclear.

One compelling brain mechanism is ‘behavioral tagging’ in which weak memories are rescued from forgetting by a subsequent stronger learning event (Ballarini et al., 2009; Moncada et al., 2011; Moncada & Viola, 2007; Wang et al., 2010). According to this model, a weak learning experience sets a ‘learning tag’ in activated synapses, creating a memory trace that is typically short-lived. However, if a stronger behavioral event engages those same neural pathways, those recent, fading traces can become stabilized in long-term memory ((see also Joels et al., 2006).

Inspired by these behavioral tagging frameworks, we used functional neuroimaging (fMRI) to determine the brain mechanisms by which emotional learning, which constitutes a strong and arousing event, selectively and retroactively prioritize memories of conceptually-related neutral information. Here, we adapted the behavioral task in Dunsmoor et al. (2015) such that participants underwent two incidental encoding phases while in the MRI scanner (Dunsmoor et al., 2015). During Phase 1 (preconditioning), participants viewed images of neutral tools and animals. Scene stimuli were also rapidly presented between these object images in an effort to provide a unique mental ‘context tag’ that was specific to information encountered Phase 1 (e.g., Gershman et al., 2013). Approximately 6 minutes later, participants then viewed a novel set of neutral tool and animal images (Phase 2). This time, one of those visual categories was probabilistically paired with a mild shock to the wrist, thereby making that conceptual information emotionally salient (Pavlovian fear conditioning). Recognition memory was then tested for all animal and tool images after a 24-hr delay.

This design allowed us to measure both ‘online’ (during the tasks) and post-encoding (‘offline’) brain mechanisms that may promote the retroactive prioritization of items related to the shocked category in Phase 2. For example, if animal images were paired with shocks during conditioning (CS+ category), we’d expect memory to be better for the animals compared to the tools from Phase 1 as well, despite neither of these stimulus categories being salient when they were first encountered. We hypothesized that one mechanism underlying this selective and retroactive memory benefit may be that neural context representations from Phase 1 are replayed or reactivated during new learning in Phase 2. This in turn would provide a transient window for emotional stimuli to selectively enhance memory for reactivated, overlapping representations. To test this idea, we used a multivoxel “scene” pattern classifier and looked for evidence of scene representations (i.e., Phase 1 context) during Phase 2 fear learning. We hypothesized that greater prior context reinstatement during conditioning would be associated with greater retroactive memory benefits for CS+ exemplars from Phase 1.

We also sought to further uncover the neuromodulatory mechanisms underlying retroactive memory by testing whether activation of the noradrenergic (NE) and dopaminergic (DA) systems during emotional learning (Phase 2) enhances the selectivity of memory consolidation. A wealth of evidence indicates that these neuromodulators promote the encoding of motivationally-relevant stimuli (Cahill et al., 1994; Lisman & Grace, 2005; Mather et al., 2015; Shohamy & Adcock, 2010; Strange et al., 2003) as well as facilitate the selective consolidation of emotional or rewarding information (Cahill & Alkire, 2003; Gruber et al., 2016; McGaugh & Roozendaal, 2002; McIntyre et al., 2002; Rossato et al., 2009; Shohamy & Adcock, 2010). Behavioral tagging work in rodents also demonstrates that NE and DA are essential for triggering the synthesis of proteins that are necessary for stabilizing weak learning experiences in long-term memory (Moncada, 2017; Moncada et al., 2011; Moncada & Viola, 2007; Wang et al., 2010). For example, in rodents, pharmacological blockade of beta-adrenoreceptors or D1/D5 can abolish the long-term mnemonic benefits of novel field exploration on weak inhibitory avoidance training (Moncada et al., 2011). We therefore hypothesized that activation of catecholaminergic nuclei (VTA/SN and LC) during emotional moments (CS+ images) might simultaneously promote mnemonic processing of overlapping past and present CS+ representations.

Finally, we considered the possibility that post-encoding consolidation processes support the selective retroactive benefit, since a study-test delay has been shown to be integral for this memory bias to occur (Dunsmoor et al., 2015; Patil et al., 2017). A fast-growing body of work demonstrates that post-encoding increases in hippocampal-cortical functional connectivity relate to behavioral measures of long-term memory (de Voogd et al., 2016; Hermans et al., 2017; Murty et al., 2016; Tambini & Davachi, 2019; Tambini et al., 2010; Tompary et al., 2015). Thus, we also tested if emotional experience-dependent increases in hippocampal connectivity with category-selective regions of visual cortex were evident and, if so, whether these changes were related to the preferential retention of Phase 1 CS+ items. Altogether, our series of fMRI analyses aimed to elucidate both online and offline neuromechansims by which emotional events selectively preserve memories that acquire salience in the future.

## Methods

### Participants

Twenty-seven healthy young adults were recruited from the New York University Psychology Subject Pool and nearby community to participate in this experiment. All participants provided written informed consent approved by the New York University Institutional Review Board and received monetary compensation for their participation. All eligible individuals were right-handed, had normal or normal-to-corrected vision and hearing, and were not taking psychoactive medications. Nine participants were excluded from data analyses for the following reasons: five participants didn’t return for session 2; two people fell asleep during scanning; one participant withdrew from session 1; and one participant had an incidental finding on his anatomical brain scan. Recruiting additional participants was not possible due to planned decommission of the MRI scanner at NYU. In total, data from eighteen participants were analyzed in this study (8 women; M_age_ = 22, SD_age_ = 2.26).

### Materials

The stimuli consisted of colored photographs used in Dunsmoor et al. (2015) as well as new scene images (Dunsmoor et al., 2015). There were three categories: 120 neutral tool images, 120 neutral animal images, and 240 outdoor scenes. Half of the outdoor scene images were phase scrambled and included in the localizer phase of the experiment. All pictures were originally obtained from the website http://www.lifeonwhite.com and from the internet. The tool and animal images were unique exemplars and had different names from each other.

### Procedure

This study involved two separate sessions that were spaced 24 hours apart. In the first session (fMRI), participants’ brains were scanned during a scene functional localizer scan, a two-phase incidental encoding task, and three intervening resting-state scans (see **Figure 1**). The resting-state scans were collected at the following times: (1) upon entering the scanner (baseline); (2) between Phase 1 and Phase 2 of the incidental encoding task (pre-conditioning); and (3) after Phase 2 of the incidental encoding task (post-conditioning). During the ∼6-m resting-state scans, participants were instructed to lay still with their eyes open while viewing a black fixation cross in the middle of the screen. An infrared camera was used to monitor pupil diameter and ensure participants did not fall asleep during these scans.

**Figure 1.**
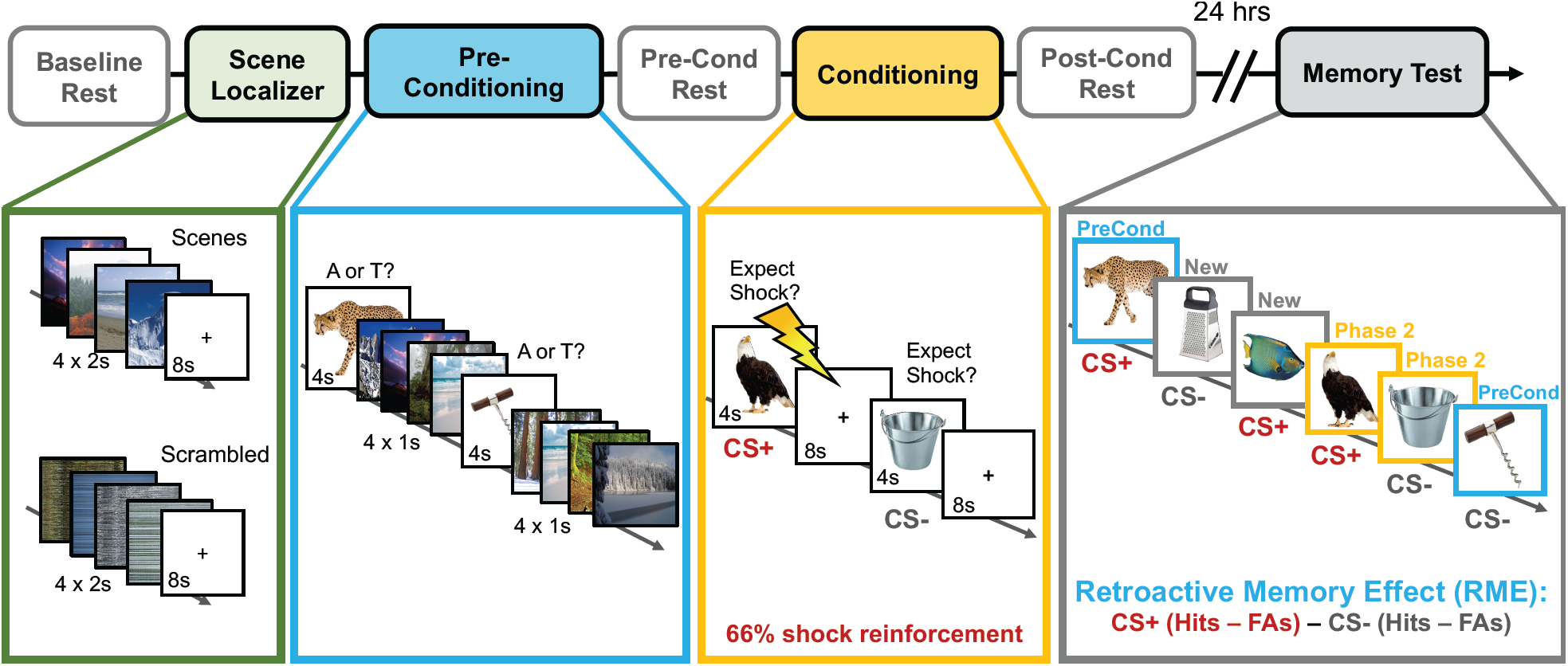
Overview of experiment design. Prior to the main two-phase incidental encoding task, participants performed a functional localizer task in which they viewed mini-blocks of neutral scenes and scrambled images (green box). Next, participants performed Phase 1 of a two-phase incidental encoding task. During this first phase (preconditioning; blue), participants viewed a series of neutral animal and tool images and had to classify the category of each image via button press. Importantly, four neutral scene images were also inserted in between each tool and animal image to create a unique ‘context tag’ for Phase 1 information. Approximately 6 minutes later, participants incidentally encoded novel animal and tool images (Phase 2; gold box). This time, however, a mild electrical shock also co-terminated with 2/3rds of the trials from one visual category (conditioning), thereby making that conceptual information emotionally significant (CS+). When each image appeared, participants rated whether they expected a shock on that trial, which provided a behavioral index of emotional learning. To examine how emotional learning influenced post-encoding hippocampal resting-state functional connectivity, resting-state scans were collected immediately before and after Phase 2 of the encoding task. Participants returned 24 hours later for a surprise recognition memory test (gray box). During this memory test, participants viewed all of the animals and tools they had seen during Phase 1 (blue borders) and Phase 2 (gold borders) of the experiment, along with new animal and tool ‘lure’ items (gray borders). Participants simply indicated whether each item had been seen previously (‘old’ judgement) or was completely new (‘new’ judgement). The critical measure in this study was the retroactive memory effect (RME), which was computed by subtracting participants’ corrected recognition memory for Phase 1 CS- items from their corrected recognition memory for Phase 1 CS+ items. Higher RME scores index a greater retroactive memory benefit for Phase 1 items that are conceptually-related to the emotional (CS+) category from Phase 2. Lightning bolt indicates shock.

**Figure 2.**
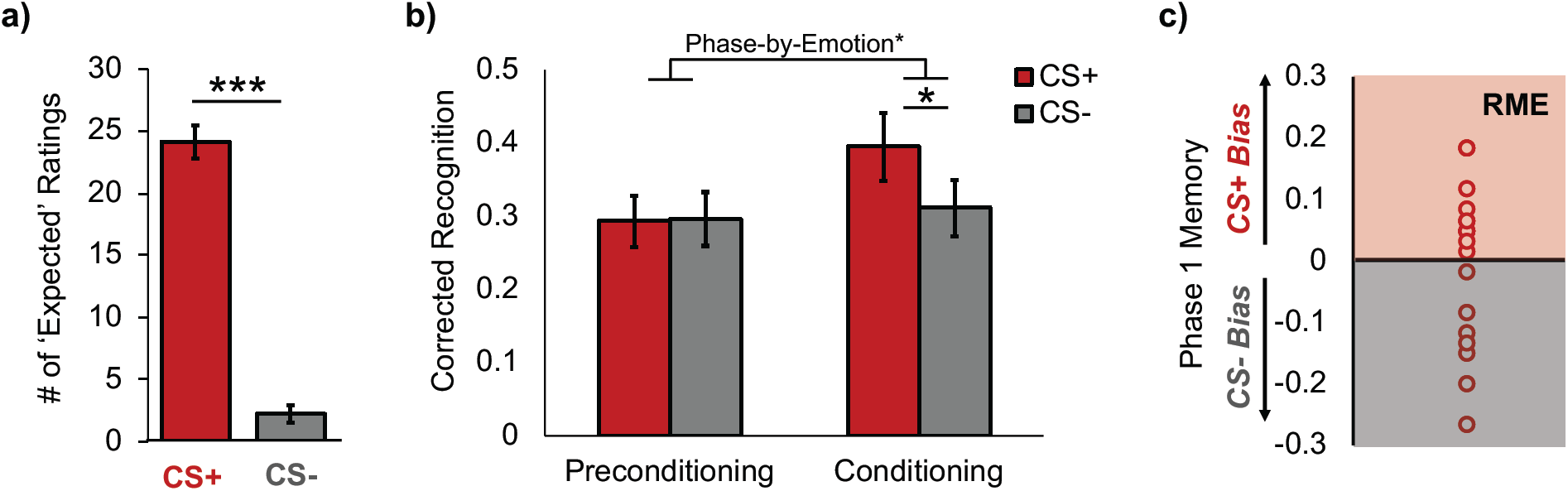
Emotional learning enhanced memory for CS+ items during conditioning, but had highly variable effects on memory selectivity for Phase 1 items. (a) Shock expectancy ratings for CS+ and CS- trials verified that participants were able to learn which visual category was paired with shock. (b) After a 24-hr delay, participants showed an emotional memory enhancement for CS+ (red bars) compared to CS- items (gray bars) from the conditioning phase of the experiment (Phase 2). However, emotional learning did not retroactively bias memory in favor of conceptually-related items from the preconditioning phase (Phase 1). Corrected recognition was computed by subtracting participants’ false alarm rates from his/her hit rates for each emotion category, separately. (c) Across-participant variability in the emotion-related selective and retroactive memory effect (RME) for Phase 1 items. A subtraction score between corrected recognition rates for CS+ minus CS- items from Phase 1 is plotted on the y-axis. Values below zero represent a memory bias towards remembering items from the CS- category (gray box). Values above zero represent a memory bias towards remembering items from the CS+ category (red box), an effect that is termed “RME”. *p < .05; ***p < .001.

### Phase 1 incidental encoding (preconditioning)

In Phase 1 of the incidental encoding task (preconditioning), participants viewed a series of neutral animal and neutral tool images for 4s each (**Figure 1**). The participants’ task was to classify each image as an animal or tool via button press. During the inter-stimulus interval (ISI), a series of four outdoor scene images were presented for 1s each. Scenes were only presented during this initial encoding phase to create a unique ‘mental context tag’ for Phase 1 information (Gershman et al., 2013). A total of 30 tools and 30 animals were presented in pseudo-randomized order, such that no more than 3 objects from the same object category appeared in a row.

### Phase 2 incidental encoding (fear conditioning)

Approximately 6 minutes after Phase 1 encoding, participants viewed a new set of 30 animal and 30 tool images (Phase 2; Pavlovian fear conditioning phase). Unlike Phase 1 of incidental encoding, however, one of these categories co-terminated with a mild electrical shock on 2/3rds of the trials (20 trials; duration = 200ms; CS+ category). Each image was presented for 4s, during which time participants had to indicate whether or not they expected to be shocked on that trial. An 8-s fixation cross centered on a gray background was inserted between each image. The object categories serving as CS+/CS- (i.e., animals or tools) were counterbalanced across participants (shock category sub-groups: N_animal_ = 7, N_tool_ = 11). To verify successful emotional learning, we performed a repeated-measures analysis of variance (rm-ANOVA) on the number of CS+ and CS- trials that the participants rated they expected a shock to occur. Emotion (CS+, CS-) was modeled as the factor of interest, with Shock Category (animal, tool) as a covariate.

### Delayed recognition test

Approximately 24 hours later, participants returned and were given a surprise recognition memory test while outside of the scanner. During this test, participants viewed all of the object images they had seen in the MRI scanner as well as 60 new animal images and 60 new tool images (lures). A total of 240 images were presented in randomized order. During this self-paced memory test, the participants rated whether each item was ‘old’ (previously presented) or ‘new’ (never seen in the scanner) according to their confidence level. There were four options: ‘definitely new’, ‘maybe new’, ‘maybe old’, or ‘definitely old’. To examine the effects of emotion on recognition memory, a 2 (Emotion: CS+, CS-) x 2 (Encoding Phase: preconditioning, conditioning) mixed ANCOVA was performed on corrected recognition scores (CRS), with Shock Category (tools, animals) as a covariate. The CRS values were computed by subtracting participants’ hit rates (correctly said ‘old’) from their false alarm rates (said ‘old’ but it was new) for each encoding phase and each category, separately.

Our primary behavioral measure of interest was the degree to which participants exhibited a selective and retroactive memory benefit for CS+ exemplars from Phase 1; that is, if participants were shocked on animals during conditioning (Phase 2), they would also show better memory for animals compared to tools from the preconditioning block (Phase 1) of the task. To compute this emotion-related memory bias measure, we subtracted participants’ corrected recognition performance for CS- exemplars from their performance on CS+ exemplars that were encountered during Phase 1. Henceforth, we will simply refer to this emotion-biased retroactive memory effect measure as “RME”.

### Skin conductance response (SCR) methods

To index autonomic arousal responses during fear conditioning, SCRs were recorded via MRI-compatible electrodes placed on participants’ right wrist and measured with a BIOPAC MP100 System (Goleta, CA). Shocks were delivered to the right wrist using pre-gelled MRI-compatible electrodes connected to a stimulator (Grass Medical Instruments). Upon entering the MRI scanner, the shock electrodes were attached to the right wrist and the shock level was calibrated to be at level deemed “highly annoying but not painful” (e.g., Dunsmoor et al., 2011). Although it was not the focus of this study, we were unable to link SCRs to the appropriate trial labels due to a programming error. Thus, SCRs were not analyzed. Importantly, however, fear conditioning success was validated by trial-by-trial shock expectancy ratings, which are considered a valid measure of human conditioning with strong face- and construct-validity (Boddez et al., 2013).

## fMRI acquisition and analyses

### MRI data acquisition

All neuroimaging data were acquired on a 3T Siemens Allegra scanner located at the Center for Brain Imaging at New York University. The visual stimuli were displayed on a mirror in front of participants’ eyes that was attached to a 32-channel matrix head coil. A high-resolution T1-weighted anatomical image (MPRAGE) was acquired to aid with functional image co-registration (slices = 176 axial; TR/TE/TI = 2500ms/3.93ms/900ms; FOV = 256mm; voxel size = 1mm^3^ isotropic; slice thickness = 1mm; bandwidth = 130Hz/Px). Functional images for the three resting-state runs (184 volumes each), scene localizer task (164 volumes), preconditioning run (244 volumes), and conditioning run (364 volumes) were acquired using the same echo-planar imaging sequence (TR/TE = 2000/15 ms, 34 interleaved slices, FOV = 102 mm; FA = 82°; voxel size = 3mm^3^ isotropic).

### Image preprocessing

Image preprocessing was performed using FSL Version 5.0.4 (FMRIB’s Software Library, www.fmrib.ox.ac.uk/fsl). The first four volumes of each functional scan were discarded for signal stabilization. Functional volumes were preprocessed by removing non-brain tissue using BET, applying spatial smoothing using a Gaussian kernel of 6mm full-width-at-half-maximum (FWHM), grand-mean intensity normalization of the 4D data set by a single multiplicative factor, and applying a high-pass temporal filter of 100s. Additionally, volumes with extreme head motion artifact were regressed from the dataset. Each participant’s denoised mean functional volume was co-registered to his/her T1-weighted high-resolution anatomical image using brain-based registration (BBR) with 7 degrees of freedom. Anatomical images were then co-registered to the 2mm isotropic MNI-152 standard-space brain using an affine registration with 12 degrees of freedom.

### Parahippocampal place area region-of-interest (ROI)

Before preconditioning, a functional localizer task was used delineate the parahippocampal place area (PPA), a cortical region in the ventral visual stream that is specialized to process to scene information (Epstein & Kanwisher, 1998). The localizer scan consisted of 32 colored scenes and 32 phase-scrambled scenes. Image presentation was divided into 16 mini-blocks lasting 16-s each. Each mini-block contained either four individual scenes or four individual scrambled images lasting 2 seconds each. These image quartets were followed by an 8-s fixation cross inter-trial-interval (ITI). Participants were instructed to press a button if one of the images repeated (1-back task). None of the scenes from the localizer task were also used in the preconditioning encoding task.

A general linear model (GLM) was fit to each participant’s localizer functional data to localize and delineate the left PPA and right PPA. The GLM included separate square wave-form regressors for the scene and scrambled image mini-blocks that were convolved with a double-gamma HRF. Whole-brain statistical parametric maps were calculated for the scene > scramble contrast using a one-sample t-test. To correct for multiple comparisons, Z-statistic images were thresholded using clusters determined by *Z* > 2.3 and a corrected cluster significance threshold of *P* = .05 (Worsley, 2001). The functionally-defined ROI masks for the left/right PPA regions were defined as 6mm spheres centered upon peak voxels in the parahippocampal gyrus within the scene > scrambled whole-brain contrast map. These spheres were then merged with the same uncorrected statistical maps thresholded at *Z* = 2.57 to only retain gray matter voxels that were selective for processing scene information. Using this approach, we were able to define the PPA for all participants (mean peak MNI coordinates across participants: Left PPA [-27 -50 -10]; Right PPA [28 -47 -11]; see **Figure 3**).

**Figure 3.**
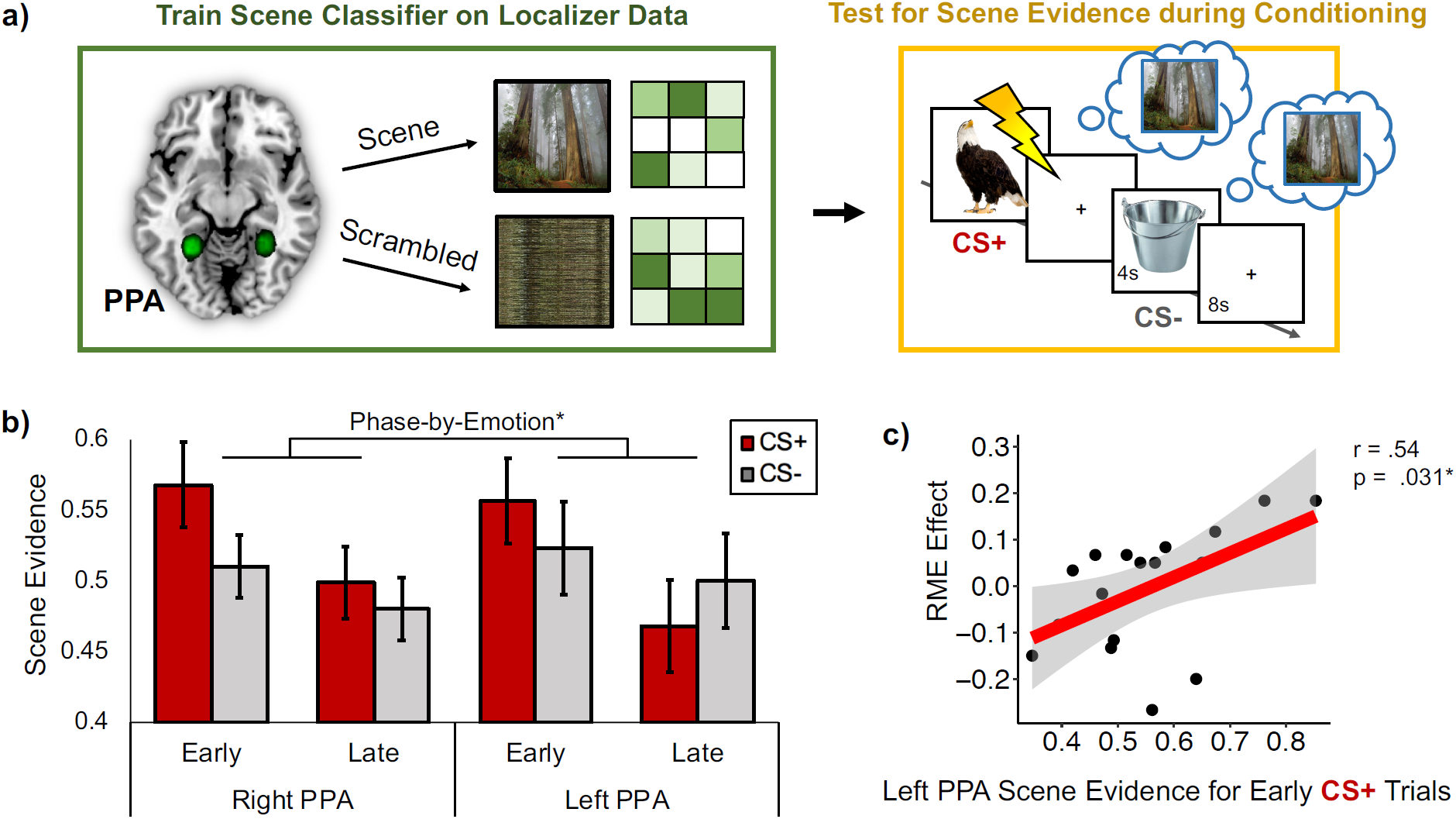
Neural reinstatement of the prior mental context increased during emotional moments in the first half of fear conditioning, and this pattern was related to greater retroactive memory benefits for items from the CS+ category. (a) A multivoxel pattern classifier was trained on the localizer data to discriminate scene versus scrambled images in the left and right parahippocampal place area (PPA; top left green box). Green circles represent study-specific probabilistic left and right PPA masks across all participants. The scene classifier was then tested on fMRI data from the conditioning phase of the experiment (Phase 2; top right gold box). Because scenes were only presented during Phase 1 of the incidental encoding task, any scene evidence output by the classifier during the CS+ and CS- images during conditioning was interpreted as reinstatement of the prior mental context (blue thought bubbles). (b) Scene evidence output by the pattern classifier was greater for emotional (CS+) compared to neutral (CS-) items during the first versus second half of conditioning. (c) A partial linear correlation analysis revealed that the emotion-related retroactive memory effect (RME) was correlated with greater scene evidence on CS+ trials from the first half of conditioning. Lightning bolt indicates shock. *p < .05.

### Regions-of-interest definitions

Target anatomical ROIs were defined for the ventral tegmental area/substantia nigra (VTA/SN) and locus coeruleus (LC), as well as animal/tool category-selective sensory cortex and left/right hippocampus. The VTA/SN anatomical mask was derived from an existing probabilistic atlas (Murty et al., 2014) and thresholded at 50% probability. A standard-space LC anatomical mask was derived from a separate study that used neuromelanin-sensitive weighted MRI to identify LC neurons in the pontine tegmentum (Keren et al., 2009).

Animal and tool-selective cortical ROIs were defined as 4mm spheres centered upon peak voxel coordinates reported in a previous fMRI study (Dunsmoor et al., 2014). The animal-selective cortical ROIs included areas of right inferior occipital gyrus and right lateral fusiform gyrus, whereas the tool-selective cortical ROIs included areas of left middle occipital gyrus and left medial fusiform gyrus. Participant-specific left and right hippocampal anatomical ROIs were extracted using the FIRST tool in FSL. These masks were then thresholded at 25% probability and binarized.

### Neural context reinstatement analysis

One of our primary goals was to determine if emotional events modulate the reactivation of prior mental contexts, and whether the degree of neural context reactivation relates to the selective consolidation of recent, emotion-related information. To test these ideas, we inserted neutral scene images between the object images in Phase 1 of the incidental encoding task (preconditioning). Because scene images were only presented during this phase of incidental encoding, they provided a unique ‘context tag’ for Phase 1 mental representations (Gershman et al., 2013). Thus, we interpret any evidence of scene information during Phase 2 as neural reinstatement of the Phase 1 mental context.

To measure the amount of scene reinstatement during emotional learning, we first trained a multivoxel pattern classifier to discriminate scenes versus scrambled scene images. Specifically, an L2-regularized multinomial logistic regression classifier was trained on multivoxel patterns of PPA BOLD signal (betas) during the scene localizer (see **Figure 3a**). An eightfold cross-validation procedure verified that the pattern classifier was highly accurate at discriminating multivoxel patterns of PPA activation between processing scene images versus scrambled scene images (mean accuracy = 96% +/ .021%).

To determine if the Phase1 mental context was reinstated during new emotional learning, we used a Least Squares All (LSA) approach to analyze the conditioning-phase data and acquire trial-by-trial estimates of multivoxel activity in the PPA. In this approach, BOLD signal for each of the conditioning-phase trials was estimated simultaneously in a single voxel-wise GLM (Rissman et al., 2004). Separate trial regressors were created by modeling the onset time for each animal and tool image with a duration of 4 seconds and convolving these regressors with a double-gamma hemodynamic response function (HRF). The six motion parameters, extreme head motion outliers, and shock deliveries (1-s stick function) were modeled as nuisance regressors in the GLM.

For each participant, individual parameter estimates were extracted separately from the left and right PPA voxels for each of the 60 conditioning-phase trials. The pattern classifier was then tested on these parameter estimates to estimate the amount of scene evidence (discrete values ranging between 0 and 1) during each conditioning-phase trial. To examine whether emotion biased the degree of Phase 1 neural context reinstatement, the classifier estimates of scene evidence were sorted by emotion trial type (CS+ or CS-).

Because accumulated evidence from Pavlovian conditioning studies suggest that amygdala and SCR signatures of fear acquisition are more robust during earlier versus later phases of fear conditioning (Buchel et al., 1998; Dunsmoor et al., 2014), we split the Phase 2 trials evenly into early and late conditioning-phase bins to see whether reinstatement of the prior context was more robust earlier on during emotional learning. A 2 (Hemisphere: left, right) x 2 (Conditioning Phase: early, late) x 2 (Emotion: CS+, CS-) mixed ANCOVA was performed on scene evidence values to examine the effects of emotion and timing on neural context reinstatement. Shock Category was modeled as a covariate.

To test our main hypothesis that online memory reactivation relates to the retroactive memory effect, we then performed partial Pearson’s correlations between participants’ RME scores and the amount of scene evidence output by the classifier for CS+ and CS- trials during conditioning, while also controlling for Shock Category. Importantly, we first conducted Shapiro-Wilk’s tests on the independent and dependent variables as well as Breusch-Pagan tests to verify that all variables used in these linear regressions were normally distributed and the correlations were homoscedastic. All of the reported regression analyses met the statistical assumptions for a Pearson’s correlation coefficient analysis.

### Emotional memory encoding GLM analysis

In addition to Phase 1 context reinstatement, we explored whether the level of engagement of neuromodulatory systems during Phase 2 fear conditioning was also related to the selective retroactive memory benefit. We reasoned that, insofar as emotion amplifies encoding processes, these enhancements may also selectively strengthen the ongoing storage of conceptually-related representations from Phase 1. To test this possibility, we first performed a subsequent memory GLM analysis for the CS+ and CS- items from Phase 2 to dissociate emotion’s specific effects on new encoding processes (see **Figure 4, left panel**). We then correlated individual differences in emotional encoding-related BOLD signal during Phase 2 to individual differences in the behavioral retroactive memory effect for Phase 1 items.

**Figure 4.**
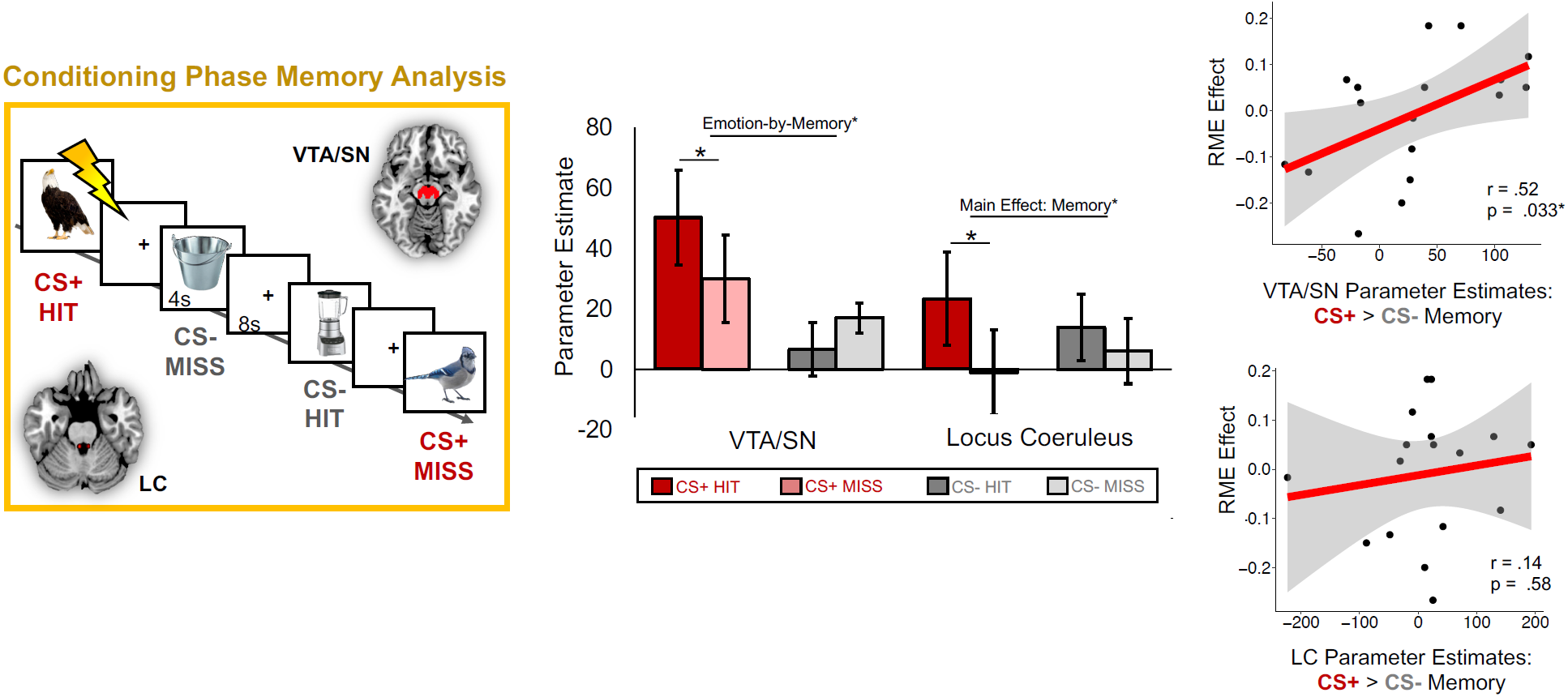
Effects of catecholaminergic nuclei activation on emotional memory, and their relationship with the retroactive memory benefit for related items. (Left Panel) A subsequent memory general linear modeling (GLM) analysis was performed for items that were incidentally encoded during the conditioning phase of the experiment. Each trial from conditioning was sorted by emotion type (CS+ or CS-) and whether it was remembered 24 hours later (hit or miss). BOLD signal was extracted from an anatomical atlas-defined VTA/SN and locus coeruleus (LC) mask for each of the conditioning-phase trials. (Middle Panel) VTA/SN activation was significantly greater when participants successfully encoded CS+ items (dark red bar), and was also more engaged during encoding of CS+ compared to CS- items during conditioning. LC activation was significantly greater during successful item encoding, which was primarily driven by memory enhancement effects for CS+ items. (Right Panel) Activation-related emotional memory enhancement scores for the VTA/SN (top) and LC (bottom) were computed by subtracting encoding-related parameter estimates for CS- trials (i.e., hit minus miss) from encoding-related parameter estimates for CS+ trials. The VTA/SN emotional memory enhancement measure was positively correlated with the magnitude of the retroactive memory benefit (RME) across participants, but LC scores were not. *p < .05.

In this Phase 2 subsequent memory GLM, we modeled separate event-related regressors for the CS+ and CS- exemplars with durations of 4-s each. Each task regressor was then convolved with a dual-gamma canonical hemodynamic response function. Next, we sorted the 60 conditioning-phase images by emotion trial-type (CS+/CS-) and by subsequent memory status (remembered: Hit; forgotten: Miss). Participant-level GLMs were constructed using 4 task regressors: (1) CS+ Hit, (2) CS+ Miss, (3) CS- Hit, and (4) CS- Miss. Additional nuisance regressors for the 6 motion parameters and extreme head movement outliers were also included in these models.

The resulting whole-brain contrast images were analyzed in higher-level mixed-effects analysis using FMRIB’s local analysis of mixed effects (FLAME 1 (Beckmann et al., 2003). A single group average for each of the contrasts-of-interest was calculated using a one-sample t-test. To correct for multiple comparisons, Z-statistic images were thresholded using clusters determined by *Z* > 2.3 and a corrected cluster significance threshold of *P* = .05 (Worsley, 2001).

To test whether neuromodulatory activity was related to emotional memory enhancements for Phase 2 items, we extracted parameter estimates for each of our four regressors-of-interest from the VTA/SN and LC. These brainstem ROI values were then submitted to separate 2 (Emotion: CS+, CS-) x 2 (Memory: hit, miss) mixed ANCOVA’s with Shock Category (shocked on animals or shocked on tools) as a covariate to examine whether activation of mesolimbic dopaminergic nuclei (VTA/SN) and noradrenergic nuclei (LC) promote the encoding of emotional information. Because there is substantial evidence implicating neuromodulators in emotional memory enhancements (e.g., McGaugh, 2013; Shohamy & Adcock, 2010), we had strong directional hypotheses and chose to use one-tailed t-tests for these analyses.

Our primary goal was to examine if neuromodulatory effects during emotional learning were also associated with the preferential consolidation of Phase 1 CS+ exemplars. To this end, we first computed separate emotional memory enhancement scores for VTA/SN and LC encoding-related BOLD signal using the following formula: [CS+ (Hit > Miss)] > [CS-(Hit > Miss)]. These brain-related Phase 2 emotional encoding scores were then linearly correlated with Phase 1 RME scores using Pearson’s correlation coefficient analyses. Importantly, participants were only included in the conditioning-phase-related analyses if they had trials for all four memory/emotion bins. One participant had no trials for CS- Miss and was therefore excluded in these brainstem-related analyses (remaining N = 17).

### Emotional learning-dependent changes in post-encoding hippocampal functional connectivity

Our next important question was whether hippocampal-cortical functional connectivity were related to the selective retroactive memory benefit. Previous fMRI work has shown that, following reward or aversive learning, the hippocampus becomes more functionally coupled with regions that process the information associated with those motivationally-significant events (de Voogd et al., 2016; Murty et al., 2016). Inspired by these studies, we predicted that emotional learning may also bias subsequent hippocampal connectivity with CS+ visual cortical regions, and that this functional coupling may help preserve memory for the now-salient items encountered in Phase 1.

Here, we performed hippocampal seed-based functional connectivity analyses. We first extracted the mean hippocampal BOLD timeseries from each participant’s three resting-state scans (see **Figure 1** for timings). These hippocampal activity timeseries were then modeled as regressors of interest in separate whole-brain GLM’s. Additional nuisance regressors for the 6 motion parameters, extreme head movement outliers, and both white matter and cerebrospinal signals were also included in these models. For the latter two signals, we used FSL FAST to acquire probabilistic white matter and CSF voxel-wise masks for each participant. These masks were thresholded at 30% tissue-type probability and then binarized prior to extracting nuisance BOLD signal.

To quantify experience-dependent changes in hippocampal functional connectivity, we subtracted hippocampal-ROI connectivity estimates from the pre-conditioning resting-state scan from connectivity estimates from the post-conditioning resting-state scan. Finally, we performed Pearson’s partial correlations to test our hypothesis that increased hippocampal connectivity with neuromodulatory nuclei and CS+ cortex after emotional learning would relate to RME scores across participants. Because we did not have explicit predictions about hippocampal laterality effects, we collapsed the ROI results across brain hemispheres.

### Mediation analysis

In the final analysis, we examined if emotion-related changes in hippocampal functional connectivity accounts for the relationship between VTA/SN emotional encoding processes and the retroactive memory effect. For this, we used the mediation package in R. The significance of this mediation model was tested using nonparametric bootstrapping with 1000 iterations. We report on the average causal mediation effect (ACME).

## Behavioral Results

### Shock expectancy ratings and recognition memory

During the conditioning phase, participants were significantly more likely to indicate that they expected a shock on CS+ compared with CS- trials, verifying successful fear acquisition at the category level, F(1,16) = 159.51, p < .001, η_p_ ^2^ = 0.91 (**Figure 2a**).

With respect to delayed recognition memory performance, there were no significant differences in false alarm rates between CS+ and CS- category lure items, F(1,16) = 0.006, p = .94, η_p_ ^2^ = 0. A 2 (Encoding Phase: preconditioning, conditioning) x 2 (Emotion: CS+, CS-) ANCOVA with Shock Category as a covariate revealed a marginally significant main effect of Encoding Phase, F(1,16) = 4.34, p = .054, η_p_ ^2^ = 0.21, on corrected recognition scores, with memory performance being better for items encoded during the conditioning compared to the preconditioning phase of encoding (**Figure 2b**). However, there was no significant main effect of Emotion on corrected recognition scores, F(1,16) = 2.00, p = .18, η_p_ ^2^ = 0.11. In addition, we observed a significant encoding phase-by-emotion interaction effect on corrected recognition scores, F(1,16) = 5.29, p = .035, η_p_ ^2^ = 0.25, such that participants were significantly better at remembering CS+ compared with CS- items during the conditioning phase (Phase 2) compared with the preconditioning phase (Phase 1).

Separate follow-up planned repeated-measures ANCOVA’s on the two encoding phases revealed that recognition was significantly better for CS+ compared with CS- category items from the conditioning phase, F(1,16) = 4.83, p = .043, η_p_ ^2^ = 0.23 (**Figure 2b**). However, memory performance did not significantly differ between CS+ and CS- exemplars from the preconditioning encoding phase, F(1,16) = 0.031, p = .86, η_p_ ^2^ = .002.

As shown in **Figure 2c**, there was substantial variability in memory performance as a function of CS type. In the subsequent fMRI analyses, we leveraged this rich variability in memory performance to examine how individual differences in RME were related to various online and offline brain measures of neural reactivation and consolidation.

## FMRl Results

### Neural context reinstatement analysis

To determine if Phase 1 neural context representations were reactivated during emotional learning, we measured the amount of scene classifier evidence that was present when participants viewed CS+ and CS- exemplars during conditioning (Phase 2). A 2 (Hemisphere: left, right) x 2 (Conditioning Phase: early, late) x 2 (Emotion: CS+, CS-) repeated-measures ANCOVA with Shock Category as a covariate revealed that there was greater scene evidence during the first compared with the second half of fear conditioning, F(1,16) = 5.81, p = .028, η_p_^2^ = .27 (**Figure 3b**). There was also a significant emotion-by-phase interaction effect, such that scene evidence was significantly greater during CS+ exemplars compared with CS- exemplars in the first versus second half of conditioning, F(1,16) = 5.46, p = .033, η_p_^2^ = 0.26. There were no other main or interaction effects on scene evidence values. These results suggest that the selective effects of emotion on prior context reinstatement is strongest during earlier versus later phases of fear conditioning

In the previous analysis, we found that scene-related neural context reinstatement was qualitatively the strongest when participants viewed emotional items during the first half of conditioning (see **Figure 3b)**. We next asked if such patterns of context reinstatement are related to the selective consolidation of conceptually-related CS+ items. A Pearson’s partial linear correlation revealed that the amount of scene evidence in left PPA on early-phase CS+ trials was significantly positively correlated with Phase 1 RME scores, partial r(15) = .52, p = .031. This brain-behavior relationship was only observed in relation to early-phase CS+ trials, as no relationship was observed with scene evidence on any other trial type (CS+ or CS-) or half of conditioning (Early or Late; all p’s > .05). Critically, these results show that neural reactivation of prior mental contexts is greater during early compared to late phases of emotional learning, and that preferential context reactivation during earlier emotional moments may serve to selectively strengthen recent, conceptually-related memories.

### Emotional memory encoding during conditioning

In the next analysis targeting ‘online’ effects of emotional learning, we examined if neuromodulatory activity was related to the encoding of CS+ exemplars during conditioning (**Figure 4, leftmost panel**). Furthermore, we were especially interested in whether emotion-related encoding patterns in the VTA/SN and LC during Phase 2 were also related to the selective and retroactive memory benefit. Of note, because we were also binning conditioning-phase trials according to memory outcome, we had low statistical power to also examine early versus late conditioning-phase effects on VTA/SN activation (as was done in the reinstatement analyses). Thus, ROI estimates were collapsed across the entirety of conditioning.

Brainstem ROI analyses revealed a significant main effect of Emotion on VTA/SN BOLD signal, F(1,16) = 5.82, p = .014, η_p_ ^2^ = 0.27, with activation being significantly greater when participants viewed CS+ items compared to CS- items during conditioning (**Figure 4, middle panel, red bar**). Planned paired t-tests revealed that VTA/SN BOLD signal was significantly higher when participants successfully encoded CS+ items, t(17) = 2.26, p = .019 (one-tailed), but not when they successfully encoded CS- items, t(16) = -0.98, p = .17 (one-tailed), during Phase 2. This pattern was qualified by a significant emotion-by-memory interaction effect, F(1,16) = 3.64, p = .033, η_p_ ^2^ = .19 (one-tailed), with CS+ trials leading to greater encoding-related VTA/SN activation than CS- trials.

For the LC, there was a significant main effect of Memory Outcome on BOLD signal, with participants exhibiting greater LC BOLD signal during successful item encoding overall, F(1,16) = 3.52, p = .040, η_p_ ^2^ = .18 (one-tailed). Separate planned follow-up t-tests on LC activation revealed a significant main effect of successful item encoding on LC BOLD signal for CS+ items, t(17) = 1.84, p = .042 (one-tailed), but not CS- items, t(16) = 0.32, p = .27 (one-tailed), during Phase 2. We did not observe any other main or interaction effects on LC activation (p’s > .05).

Importantly, across participants, the magnitude of the Phase 2 emotional memory enhancement supported by VT/SN activation was significantly positively correlated with the extent of RME across participants, partial r(17) = .52, p = .033 (**Figure 4, rightmost panel**). By contrast, there was no significant correlation between LC emotional memory activation and RME scores, partial r(17) = .14, p = .58. These findings suggest that dopaminergic activity not only selectively promotes encoding of emotional material but also enhances the selective consolidation of recently encoded related information. Thus, the local effects of emotion on VTA/SN encoding processes also appear to converge with the selective and ongoing consolidation of overlapping memories. While LC activation was associated with enhanced memory encoding, especially for emotion-related items, this modulation was not associated with the retroactive memory effect.

### Post-encoding hippocampal functional connectivity results

In the next GLM analysis, we examined whether emotional learning leads to biases in post-encoding hippocampal functional connectivity. Four separate 2 (Hemisphere: left hippocampus, right hippocampus) x 2 (Rest Phase: preconditioning rest, postconditioning rest) ANCOVAs with Shock Category as a covariate revealed that, on average, emotional learning did not significantly alter hippocampal functional connectivity with the LC, VTA/SN, CS- cortex, or CS+ cortex (all p’s > .05).

While we did not observe any main carryover effects of emotional learning on overall hippocampal connectivity, we were primarily interested whether variability in hippocampal functional connectivity changes was related to the magnitude of RME effects. In these linear correlation analyses, hippocampal pre-to-post emotional learning functional connectivity values were collapsed across hemispheres, given we did not observe any interactions between these values and brain hemisphere in the prior analysis.

Consistent with our main hypotheses, Phase 1 RME scores were indeed positively correlated with increased hippocampal-CS+ category-selective cortex functional connectivity, partial r(15) = .62, p = .0082, but not with hippocampal-CS- category-selective cortex functional connectivity, partial r(15) = -.13, p = .61, following emotional learning (**Figure 5**). A Williams test for dependent correlations indicated that these two brain-behavior correlations were also significantly different from each other, t = -2.95, p < .01. This finding suggests that emotional learning may enhance the selective consolidation of conceptually-related stimuli by biasing post-encoding hippocampal connectivity to target sensory regions associated with the emotional stimuli. When examining connectivity patterns with brainstem nuclei, we did not observe any significant correlations between hippocampal changes in functional coupling with the LC, partial r(15) = .12, p = .64, or VTA/SN, partial r(15) = .34, p = .18.

**Figure 5.**
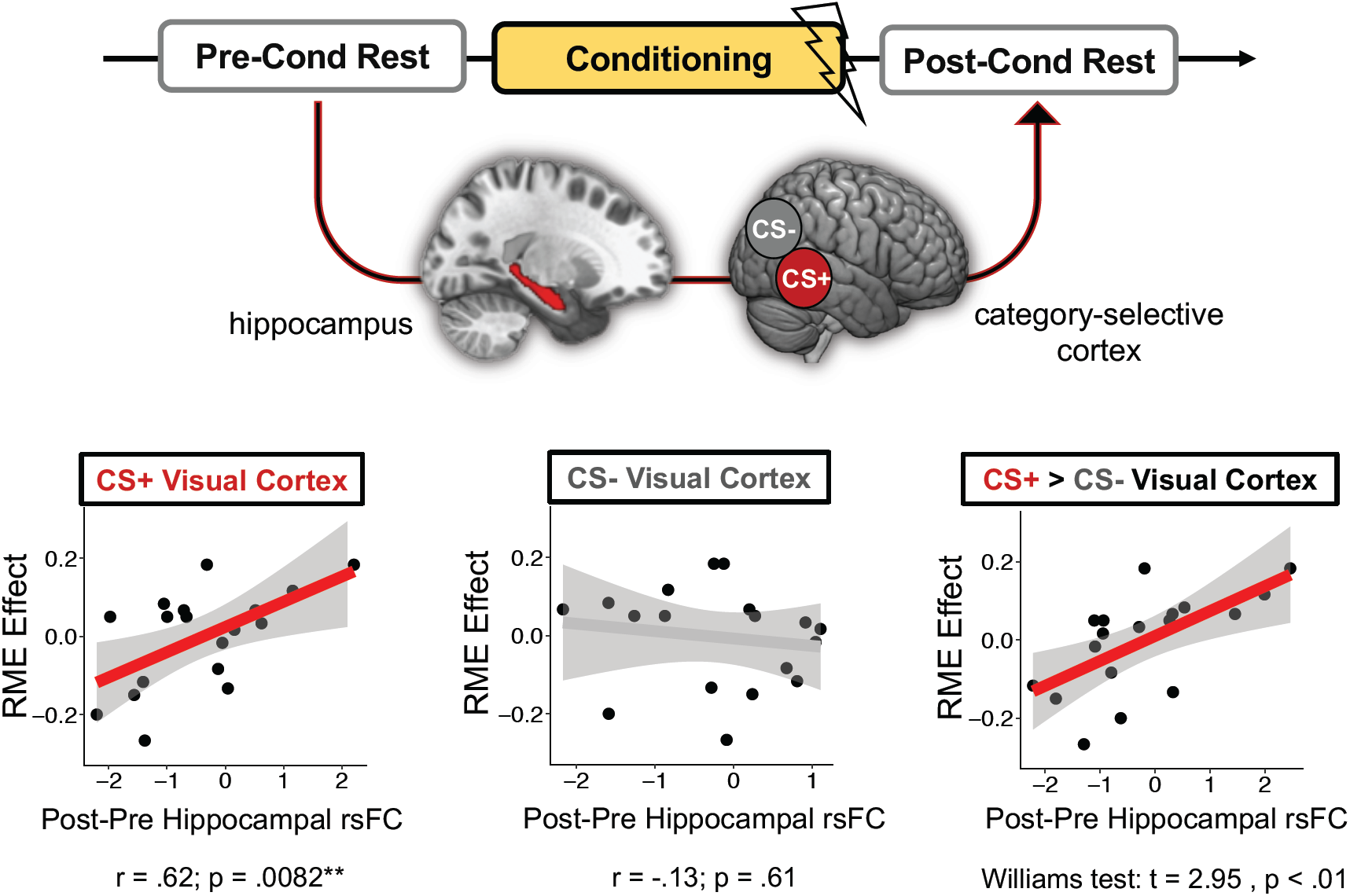
functional connectivity relate to across-participant variability in the retroactive memory effect (RME). Conditioning-dependent changes in hippocampal resting-state functional connectivity (rsFC) with CS+ (emotional) and CS- category-selective cortex were assessed by subtracting connectivity z-stats for pre-conditioning rest from connectivity z-stats from post-conditioning rest. Across participants, greater emotion-related RME effects were associated with greater hippocampal functional coupling with CS+ category-selective cortex (red line) but not CS- category-selective cortex (gray line). These hippocampal functional connectivity patterns were also significantly different from each other, indicating that, following emotional learning, greater selective retroactive memory benefits relate to a shift in hippocampal coupling towards CS+ cortex and away from CS- cortex.

To ensure that these brain-behavior relationships were specifically driven by emotional learning, we performed additional control analyses. For these regressions, we queried hippocampal-cortical functional connectivity changes from baseline rest period to the rest period following Phase 1 encoding (preconditioning; see **Figure 1**). Because neither category of information was differentially salient during Phase 1 encoding, we did not expect there to be any relationships between hippocampal connectivity patterns and RME scores. Indeed, the control analyses revealed no significant relationships between RME scores and changes in hippocampal functional coupling with CS- category-selective cortex, partial r(15) = -.36, p = .15, CS+ cortex, partial r(15) = -.40, p = .11, or the difference between the two (Williams test: t = 0.11, p < .91). Furthermore, there were no significant correlations between RME scores and hippocampal functional connectivity changes with the LC, partial r(15) = .019, p = .94, or VTA/SN, partial r(15) = -.34, p = .18. These analyses help verify that memory-related biases in hippocampal connectivity were indeed experience-dependent.

## Emotional learning-dependent changes in post-encoding hippocampal Mediation analysis

So far, the results have revealed that both online (conditioning-phase VTA/SN activation and classifier evidence during CS+) and offline (pre-to-post conditioning hippocampal-cortical functional connectivity) brain measures relate to the retroactive memory effect. In the next analysis, we examined whether these brain measures were also correlated each other. Pearson’s correlation coefficient analyses revealed a significant correlation between VTA/SN emotion-related encoding activation and pre-to-post learning changes in hippocampal-CS+ category-selective cortex functional connectivity, partial r(14) = .50, p = .048. By contrast, the degree of early neural context reinstatement in left PPA on CS+ trials (when reinstatement was qualitatively strongest) did not correlate with either of these measures (p’s > .05). These findings provided initial evidence that the online effects of dopaminergic activity may interact with subsequent consolidation processes to influence memory selectivity.

To explore this relationship further, we examined if these emotion-related biases in post-encoding hippocampal-cortical functional connectivity could account for the association between online VTA/SN encoding-related activation and RME behavioral scores. Consistent with this possibility, we found that, across participants, increased pre-to-post emotional learning changes in hippocampal-CS+ category-selective cortex connectivity mediated the relationship between Phase 2 VTA/SN emotional encoding effects and Phase 1 RME scores (ACME = 0.00047; p = .042; **Figure 6**).

**Figure 6.**
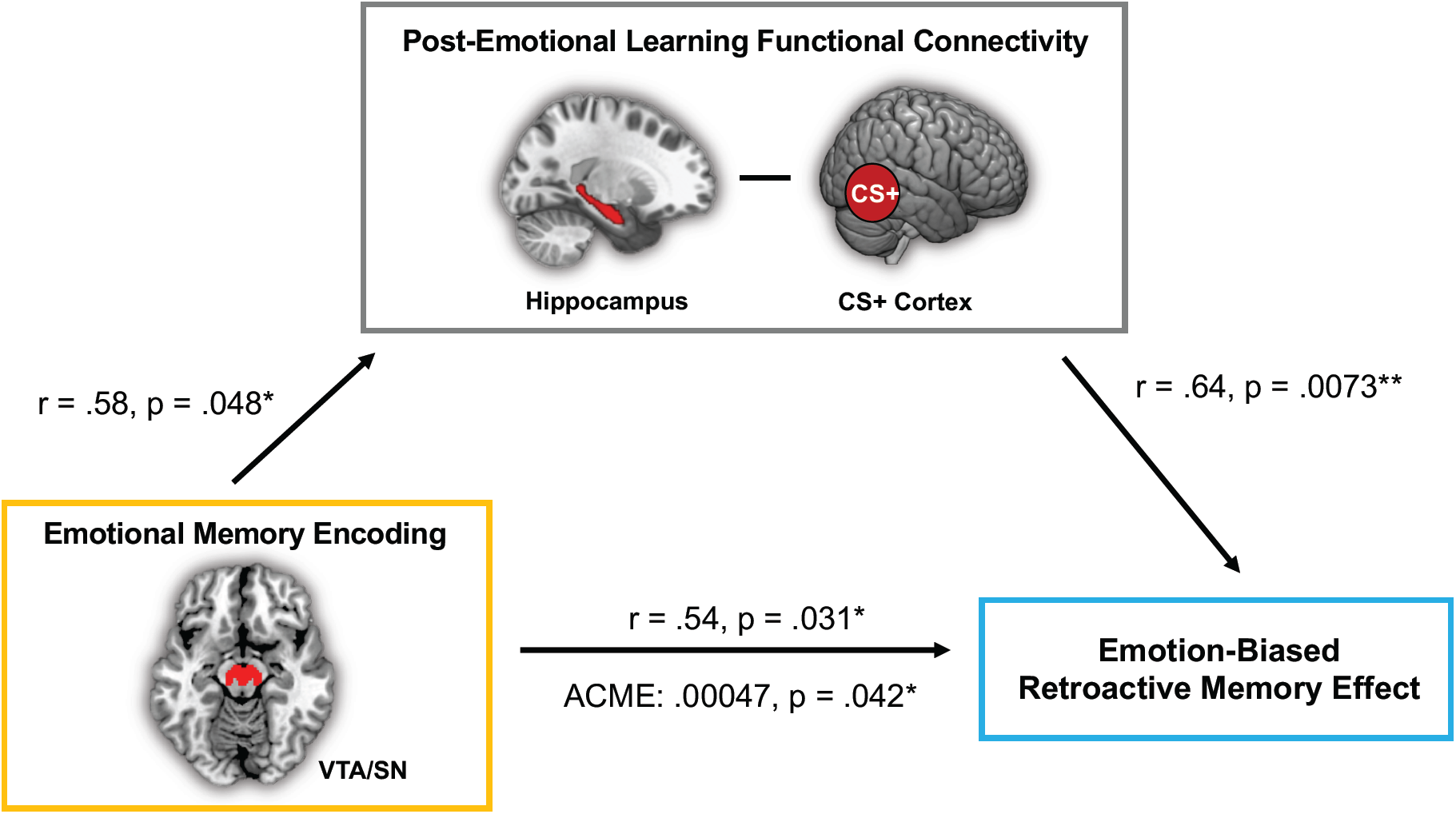
Post-encoding hippocampal functional connectivity with CS+ category-selective cortex mediates the relationship between VTA/SN emotional encoding activation and the retroactive memory benefit. Path values represent partial Pearson’s correlation coefficients after controlling for the effects of Shock Category (shocked on animals or shocked on tools). *p < .05; **p < .01.

To determine the specificity of this mediation effect to Phase 1 memory biases, we also performed the same correlation analyses, but this time targeting emotional memory biases for Phase 2 items. This memory measure (like RME) was computed by subtracting corrected recognition scores between Phase 2 CS+ and CS- items. In contrast to the brain-behavior relationships identified for Phase 1 memory, we did not find any significant associations between Phase 2 emotion-related memory biases and either the VTA/SN effect, partial r(14) = .29, p = .28, or hippocampal-CS+ connectivity, partial r(15) = .28, p = .27, across participants. Moreover, the same type of mediation analysis as before was not significant (ACME = .00011, p = .66).

Together, these findings suggest that offline hippocampal processes may stabilize recent dopaminergic learning ‘tags’ for salient concepts. Consistent with models of behavioral tagging, this consolidation process may only be necessary for preserving weaker memories of more distantly encountered, conceptually-related items. By contrast, emotional representations (i.e., those items paired with shock) may be sufficiently enhanced during initial encoding, as encoding-related VTA/SN activation was related to better memory for CS+ compared to CS- images.

## Discussion

Decades of research show that emotional experiences are both vividly and enduringly remembered. Yet recent evidence suggests that emotional arousal can also selectively influence the consolidation of more distantly encountered neutral information so long as it is conceptually-related to the emotional event (Dunsmoor et al., 2015). Here, we demonstrate that across-participant variability in emotional tagging processes relate to increases in: (1) reactivation of a prior mental context during the initial phases of new emotional learning; (2) encoding-related VTA/SN activation for emotional versus compared to neutral items; and (3) emotional learning-dependent changes in post-encoding hippocampal functional connectivity with cortical regions associated with the emotional stimulus. Importantly, post-encoding hippocampal-cortical functional coupling mediated the relationship between dopamine-related emotional memory enhancements and the retroactive memory benefit. This finding suggests that post-encoding hippocampal-cortical interactions may promote the stabilization of dopaminergic ‘relevance tags’ in category-selective cortex, providing a mechanism for adaptively storing recent, seemingly mundane information in long-term memory.

Building on a series of experiments in rodents, the behavioral tagging model posits that a weak memory trace can be strengthened by a stronger, arousing event that engages the same neural pathways (Ballarini et al., 2009; Moncada & Viola, 2007). Inspired by this idea, we interspersed neutral scene images between animal and tool images to create a unique ‘context tag’ for this block of encoding. We then trained a multivoxel “scene” pattern classifier to determine if those prior representations and neural pathways become reactivated, or re-engaged, during Pavlovian category-specific fear conditioning several minutes later. We found that Phase 1 context reinstatement, as indexed by classifier evidence of scene-related processing, was significantly more likely to occur during emotionally-significant images (CS+) than mundane images (CS-) during the first versus the second half of fear conditioning. Further, the amount of prior context reactivation at these moments was also associated with the extent of the retroactive memory benefit. Converging findings in rodents and humans demonstrate that the reinstatement of existing memories during new learning can influence how they are stored and/or updated over time (Gershman et al., 2013; Girardeau et al., 2017; Kuhl et al., 2010; Tambini & Davachi, 2019). For instance, the degree of neural item reinstatement during reward learning has been shown to increase implicit preferences for those neutral items, even though they had never been directly associated with reward (Wimmer & Shohamy, 2012). Here, we expand upon these findings by showing that emotional events may promote neural reinstatement processes of prior neutral experiences, which also has consequences for how those reactivated memory representations are stored for the long term.

Interestingly, our pattern classification results also suggest that neural context reactivation is most evident during the earlier phases of emotional learning. Previous studies using Pavlovian conditioning show that amygdala and SCR indices of fear responses tend to be most evident during the earlier portions of traditional fear conditioning (Buchel et al., 1998; LaBar et al., 1998) and tool/animal category-conditioning procedures (Dunsmoor et al., 2014). Thus, one intriguing possibility is that the spread of emotional significance to related memory representations is greatest when emotional events are most novel, salient, and arousing at the beginning of conditioning.

A core feature of the behavioral tagging model is that DA and NE release are necessary for triggering the production of proteins that can transform weak learning tags into more enduring memory traces (Moncada, 2017; Moncada et al., 2011; Ritchey et al., 2016; Wang et al., 2010). Earlier work demonstrating that these neuromodulators can convert early-phase long-term potentiation (LTP) processes to a more persistent form of late-phase LTP laid important groundwork for experiments targeting memory expression (Frey & Morris, 1997; Straube et al., 2003). In addition to modulating consolidation processes, much work also shows that dopaminergic and noradrenergic activity promote the encoding of motivationally-relevant information in long-term memory (Cahill et al., 1994; O’Carroll et al., 2006; Rossato et al., 2009; Shohamy & Adcock, 2010; Strange et al., 2003). Taken together, these converging lines of work suggest that emotional moments activate catecholaminergic systems, and, by extension, facilitate the strongest modulation of synaptic plasticity for activated synapses. Aligning with this idea, we found that greater VTA/SN activation during encoding of emotional versus neutral events also relates to selective consolidation of conceptually-related stimuli encountered several minutes earlier. Thus, strong dopaminergic activity during the encoding of CS+ exemplars may have selectively triggered consolidation processes in synaptic pathways related to and associated with the relevant emotional information.

In addition to finding ‘online’ effects of emotional learning on consolidation, we also found that individuals who showed greater hippocampal functional coupling with CS+ category-selective cortex after emotional learning also showed a larger memory benefit for CS+ category items from Phase 1 of encoding. This finding adds to a growing literature implicating post-encoding hippocampal-cortical functional coupling in the preferential retention of motivationally-significant memories (de Voogd et al., 2016; Murty et al., 2016). Our data expand upon this work by showing that post-encoding hippocampal connectivity may also be strengthening more remote representations encoded in the same neural pathways. Interestingly, we also found that these increases in hippocampal coupling with CS+ category-selective cortex mediated the relationship between online dopaminergic emotional encoding processes and the retroactive memory benefit. The long-term benefits of dopaminergic activity on memory have been previously linked to cellular consolidation mechanisms, including the stabilization of hippocampal plasticity (Lisman et al., 2011; Lisman & Grace, 2005) as well as persistent memory reactivation in hippocampal neurons (McNamara et al., 2014). In the same vein, hippocampal-cortical interactions are thought to selectively facilitate the storage of recent information that received a ‘salience’ or ‘behavioral’ tag at encoding (Moncada et al., 2015; Wang et al., 2010) Here, we provide evidence in humans that dopaminergic processes may provide such a ‘relevance tag’ for motivationally-relevant information, which is then stabilized by hippocampal processes to promote the preferential retention of weakly encoded but related memories.

Although our findings shed new light on how emotion influences the selectivity of memory consolidation, there are several limitations that warrant consideration. First, we had a modest number of participants, so additional work will be necessary to replicate and validate these effects. Second, in contrast to earlier work (Dunsmoor et al., 2015; Patil et al., 2017), we did not find a significant emotion-related retroactive memory benefit in behavior. One reason for the null effect in our study may have been the introduction of neutral scene stimuli during the initial encoding phase, which may have distracted individuals from encoding the animal and tool stimuli. Third, due to equipment malfunction, we were unable to collect skin conductance measures as an endogenous index of fear acquisition. We do not believe, however, that this detracts from our interpretations concerning emotion or salience-related effects on selective consolidation. In the current design, we weren’t specifically interested in evaluating the efficacy of fear conditioning but rather how imbuing existing neutral memories with motivational significance affects their long-term consolidation. Thus, we were able to verify that participants learned the emotional significance of the stimuli using their shock expectancy ratings, which are considered a valid measure of human fear conditioning with strong face- and construct-validity (Boddez et al., 2013).

Another possibility for the null memory effect is that being in the MRI increased participants’ baseline arousal to different degrees, which may have overshadowed the retroactive effects of the fear conditioning manipulation on Phase 1 encoding (Muehlhan et al., 2011). Despite not having measures of autonomic activation, however, we found that activation of arousal-related neuromodulatory systems (i.e., DA) was significantly higher for CS+ compared with CS- items during emotional learning, suggesting that CS+ images were indeed salient and processed as motivationally relevant (e.g., (Shohamy & Adcock, 2010). Disentangling whether these DA effects – and retroactive memory effects more broadly – relate to stimulus salience, emotionality, attention, or physiological arousal is an important direction for future research.

The current findings may also inspire future studies geared towards understanding the specific conditions under which emotion’s modulatory effects will spread to recent memories. For instance, existing theories posit that strong, emotionally arousing events will selectively enhance learning for information that engages a common neural substrate and is encountered close by in time (Ballarini et al., 2009; Joels et al., 2006; Moncada & Viola, 2007). Thus, while we have focused on the spatial (neural overlap) convergence between two learning events in the present study, the temporal proximity of these events is also important for linking them together in memory. Indeed, recent work in rodents demonstrates that an aversive event will only become associated in memory with a recent, weaker learning event if they occur in close temporal proximity to each other (i.e., less than 6 hours apart; Cai et al., 2016; Rashid et al., 2016).

It will also be important to identify any boundary conditions for emotion’s retroactive influence on recent memories; that is, determining the extent to which emotional and neutral stimuli must overlap (and how) for there to be an effect. Emotion generalization can be driven by the amount of conceptual or perceptual resemblance between an affective stimulus and other neutral stimuli (Dunsmoor & Murphy, 2014; Verosky & Todorov, 2010). It has also been shown that post-encoding stress only modulates memory consolidation when it is administered in the same room as encoding, suggesting that emotion’s and/or physiological arousal’s retroactive effects on memory may also be constrained by the learning context and not just the overlap between the target memoranda (Sazma, McCullough, et al., 2019; Sazma, Shields, et al., 2019).

In summary, our findings reveal key evidence that multiple online and offline neural processes help to adaptively prioritize the consolidation of both salient and seemingly mundane information. Acquiring a better characterization of the factors that engage these memory mechanisms is central to understanding how aversive associations may spread and persist in post-traumatic stress disorder (PTSD) and phobias. At the same time, future studies may also inform how manipulating the conceptual or contextual overlap between to-be-encoded material and a stimulating event can be leveraged to benefit new learning.

## Acknowledgements

The authors thank Darren Yi for his assistance with data collection and programming. This project was funded by federal NIH grant R01 MH074692 to L.D. and by a fellowship on federal NIH grants T32 MH019524 and F32 MH114536 to D.C.

